# Oligomerization of NLR immune receptor RPP7 triggered by atypical resistance protein RPW8/HR as ligand

**DOI:** 10.1101/682807

**Authors:** Lei Li, Anette Habring, Kai Wang, Detlef Weigel

## Abstract

In certain plant hybrids, autoimmunity is triggered by immune components that interact in the absence of a pathogen trigger. Often, NLR immune receptors are involved, with a particularly interesting case in *Arabidopsis thaliana* involving variants of the NLR RPP7 as well as variants of RPW8/HR proteins, which are homologs of animal MLKL and fungal HELL domain proteins. We demonstrate that HR4^Fei-0^ but not the closely related HR4^Col-0^ protein directly disrupts intramolecular association of RPP7b^Lerik1-3^, which in turn initiates P-loop dependent NLR signaling. In agreement, RPP7b^Lerik1-3^ forms a higher-order complex only in the presence of HR4^Fei-0^ but not HR4^Col-0^. In addition, we find that HR4^Fei-0^ on its own can form detergent-resistant oligomers suggestive of amyloid-like aggregates, which in turn can directly kill cells in an RPP7b^Lerik1-3^-independent manner. Our work provides in vivo biochemical evidence for a plant resistosome complex and the mechanisms by which RPW8/HR proteins trigger cell death.

## INTRODUCTION

Multicellular organisms and their microbial pathogens are engaged in a perpetual evolutionary race, and the genes hosts use to stay ahead of their enemies generally belong to the most variable and diverse gene families. This is the case for the dominant type of intracellular plant immune receptors, nucleotide-binding and leucine rich repeat domain (NLR) proteins, which are closely related to animal NOD/CARD proteins (Jones et al., 2016; Maekawa et al., 2011a). Plant NLRs can be classified into CC-NLRs (CNLs) and TIR-NLRs (TNLs) based on their N-termini, which include either coiled-coil (CC) or Toll/interleukin-1 receptor (TIR) domains (Qi and Innes, 2013; Zhang et al., 2017a).

Plant NLRs directly or indirectly recognize non-self molecules mainly, though not exclusively, through their C-terminal LRR domain (Ade et al., 2007; Dodds et al., 2006; El Kasmi et al., 2017; Krasileva et al., 2010; Rairdan et al., 2008; Steinbrenner et al., 2015). The detection of non-self molecules leads to conformational changes, followed by the release of intramolecular interactions, which in turn expose the N-terminal CC or TIR domains for participation in downstream signaling (Bernoux et al., 2011; Casey et al., 2016; Cesari et al., 2016; Maekawa et al., 2011b; Wang et al., 2019a, 2019b).

The importance of self-associations for the function of plant NLR proteins has been highlighted in several studies. For example, the TNLs SNC1, N and RPP1 can all form oligomers, or at least dimers, either before or after activation (Mestre and Baulcombe, 2006; Schreiber et al., 2016; Xu et al., 2014). Two different interfaces in the TIR domains of SNC1 and RPP1 can support self-association, potentially enabling TNLs to assemble higher-order complexes (Zhang et al., 2017b). Similarly, several CNL full-length receptors, such as MLA, Sr33, Sr50, RPS5 and Rx, can self-associate (Ade et al., 2007; Casey et al., 2016; Cesari et al., 2016; Maekawa et al., 2011b; Moffett et al., 2002). In addition, structural and phenotypic analyses have indicated that an extended CC fragment of Sr33 can dimerize and activate signaling, while a shorter fragment cannot (Casey et al., 2016). Cryo-EM analysis of the CNL ZAR1 has revealed how uridylation of the ZAR1 ligand PBL2 induces ZAR1 oligomerization in vitro. In its resting state, ADP-bound ZAR1 is complexed with adaptor protein RKS1 in a conformation that buries the oligomerization interfaces. Upon interaction of RKS1 with uridylated effector PBL2, ADP release allows ATP/dATP to enter the binding pocket, triggering a ZAR1 conformational change that exposes the oligomerization interfaces, which finally leads to formation of a wheel-like pentameric complex. A key event during oligomerization is the rearrangement of the CC domains, which end up forming a funnel-shaped structure that has been speculated to directly disrupt the plasma membrane (Dangl and Jones, 2019; Wang et al., 2019a, 2019b).

Some NLRs acts as sensors that positively regulate helper NLRs, although it is unknown whether these helper and sensor NLRs reside in a single complex. For example, tobacco NRG1, *A. thaliana* ADR1, and Solanaceae NRCs mediate signaling activated by multiple plant sensor NLRs (Bonardi et al., 2011; Castel et al., 2018; Collier et al., 2011; Peart et al., 2005; Wu et al., 2017, 2018b), likely interacting only indirectly with TNLs (Qi et al., 2018). Notably, these helper NLRs belong to a subclass of CNLs whose CC domains are similar to parts of the non-NLR RPW8/HR immune proteins (and therefore referred to as CC_R_ domains) (Collier et al., 2011). Similar to many NLR genes, the CC_R_-only *RPW8*/*HR* genes are located in a gene cluster, with extensive copy number and sequence variation between wild strains (Barragan et al., 2019; Jorgensen and Emerson, 2008; Orgil et al., 2007; Xiao et al., 2004). However, different from typical NLRs, which function primarily in race-specific resistance, *RPW8*/*HR* genes seem to be mostly involved in broad spectrum resistance (Ma et al., 2014; Wang et al., 2007; Xiao et al., 2001). The CC_R_ domain is predicted to be structurally related to portions of animal mixed-lineage kinase domain-like (MLKL) and fungal HeLo and HeLo-like (HELL) domains, which share structures characterized by four-helix bundles (Bentham et al., 2018; Jubic et al., 2019). Similar to the fungal HeLo domain protein Het-S, RPW8/HR is predicted to have an N-terminal transmembrane domain, a central CC_R_ /MLKL/HELL domain, as well as C-terminal repeats (Barragan et al., 2019). The N-terminal domain of Het-S can insert into membranes, thereby forming a pore and inducing cell death, while the regulatory C-terminal 21-amino acid repeats constitute a prion-forming domain (PFD) that can regulate Het-S activity by adopting an amyloidal structure (Daskalov et al., 2016; Greenwald et al., 2010; Seuring et al., 2012).

There has been no prior evidence that RPW8/HR proteins function as helpers to sensor NLRs, although genetic evidence has linked different alleles of the CNL locus *RPP7* to different alleles at the *RPW8/HR* locus. When wild strains of *A. thaliana* are crossed, the F_1_ hybrid progeny of specific crosses can express autoimmunity indicative of activated NLR signaling in the absence of a pathogen trigger. In at least three such cases, the causal factors are a specific allele of *RPP7* from one parent and a specific allele of *RPW8/HR* from another parent (Barragan et al., 2019; Chae et al., 2014). Here, we demonstrate that RPW8/HR proteins can directly interact in an allele-specific manner with RPP7 proteins. The interaction disrupts RPP7 intramolecular association, which in turn leads to assembly of a higher-order RPP7 complex and autoimmune signaling; these findings provide biochemical evidence for the in vivo formation of a plant resistosome complex, which before only has been observed in vitro (Wang et al., 2019a, 2019b). In addition, we demonstrate that RPW8/HR proteins can kill cells on their own by forming potentially amyloid-like aggregates. Our study highlights how two different arms of plant immunity converge in form of the atypical resistance protein RPW8/HR and the conventional immune receptor RPP7.

## RESULTS

### Specificity in genetic interaction between *RPP7* and *RPW8/HR*

Our previous studies identified *RPW8.1* from KZ10 and *HR4* from Fei-0 as causal for hybrid necrosis when combined with specific alleles at the *RPP7* locus from Mrk-0 and Lerik1-3, respectively (Barragan et al., 2019). We demonstrated causality of *RPP7*^Mrk-0^ using transformation of a genomic *RPP7*^Mrk-0^*-3xFLAG* construct into the KZ10 accession and with *rpp7* frameshift mutations in Mrk-0, and we confirmed that *RPP7*^Mrk-0^ and *RPW8.1*^KZ10^ are sufficient for inducing a strong hypersensitive response in a heterologous system, *Nicotiana benthamiana* (Figure S1A, S1B, S1C).

The *RPP7* genomic locus can be complex (Guo et al., 2011), but this is not the case in Mrk-0, which has only a single *RPP7* paralog (Figure S1D). We found a more interesting situation in Lerik1-3, which similar to the reference accession Col-0 contains three paralogs, which we named *RPP7* (the ortholog of the functionally defined *RPP7* cluster member in Col-0 (Eulgem et al., 2007)), *RPP7a* and *RPP7b* (Figure S1D). Only *RPP7b* but not *RPP7* and *RPP7a* triggered a confluent hypersensitive response in combination with *HR4*^Fei-0^ in *N. benthamiana* (Figure 1A), even though RPP7b-Myc protein accumulated to much lower levels than RPP7a-Myc (Figure S1E). To further confirm *RPP7b*^Lerik1-3^ (hereafter referred to as *RPP7b*) as causal, we introduced each of the three *RPP7* paralogs, driven by the same *RPP7b* promoter, into the incompatible accession Fei-0 and the neutral Col-0 background. All *RPP7b* T_1_ plants died immediately after germination or reached at most the two-cotyledon stage, while all *RPP7* and *RPP7a* T_1_ plants appeared normal (Figure 1B). Few defects were observed in Col-0 transgenic plants (Figure S1F), with occasional defects such as curled leaves that are also seen in other NLR mis-expressers (Holt et al., 2005). Finally, we combined *RPP7b-3*×*FLAG* and *HR4*^Fei-0^*-3*×*HA* under control of their native promoters in one construct. All three independent stable T_2_ lines in Col-0 recapitulated the Lerik1-3 x Fei-0 F_2_ hybrid necrosis phenotype at 23°C (Figure 1C and S1G). Together, these results demonstrated that a specific member of the *RPP7* cluster in Lerik1-3, *RPP7b*, is the causal gene for hybrid necrosis in the Lerik1-3 × Fei-0 cross. The three *RPP7* paralogs in Lerik1-3 represent three major clades of *RPP7*-like genes in *A. thaliana*, providing a natural platform for understanding specificity in the genetic interaction between *HR4*^Fei-0^ and *RPP7b*.

**Figure 1.**
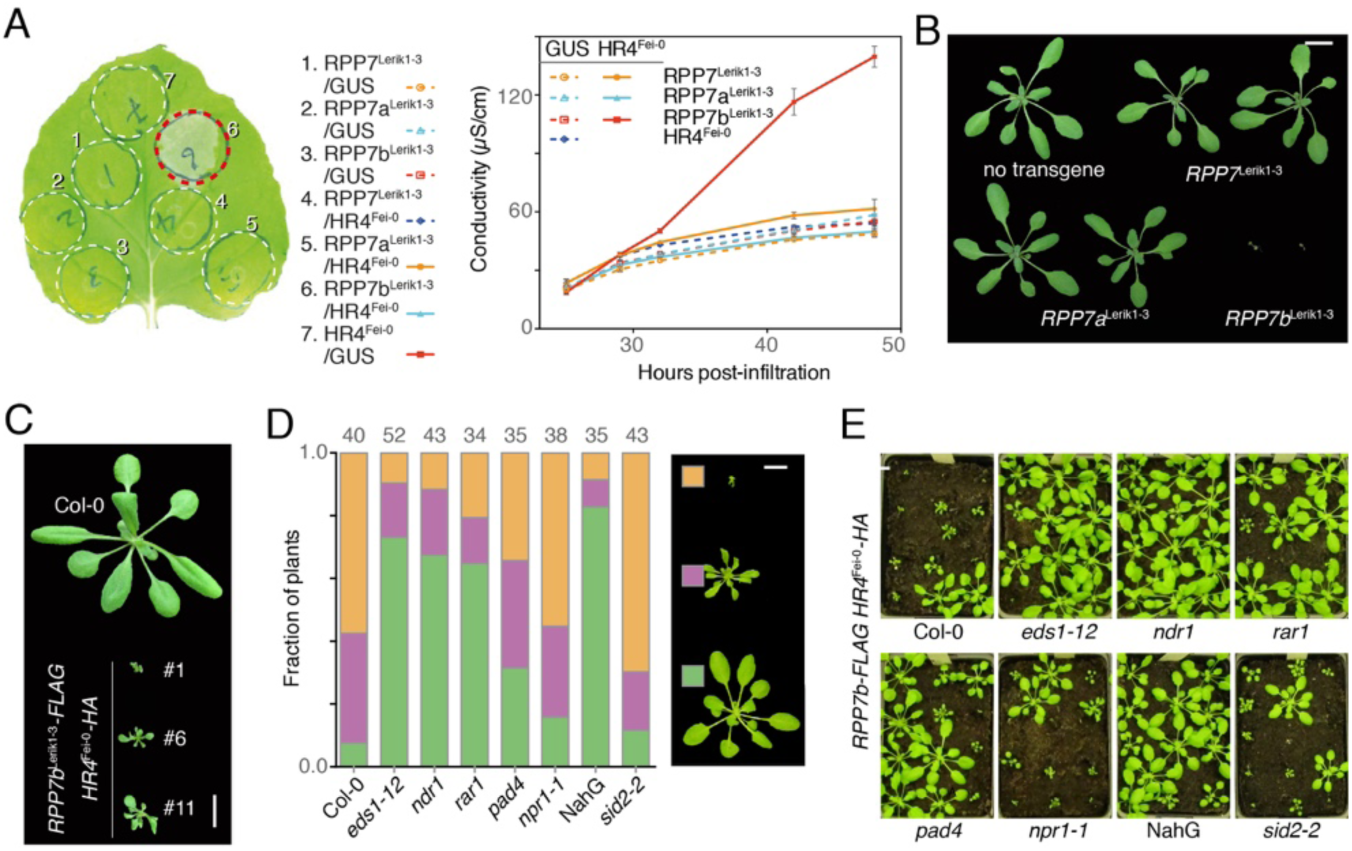
Genetics of HR4^Fei-0^/RPP7b-induced hybrid necrosis. (A) Left, hypersensitive response (indicated by a red circle) induced by co-expression of RPP7b^Lerik1-3^ with HR4^Fei-0^ in *N. benthamiana*, four days after *Agrobacterium* infiltration. Experiments were performed at least three times. Right, Ion leakage measurements of plants shown on left. Values are means ± SEM (n=3). Each sample contains eight leaf discs from different plants, and experiments were performed at least twice. (B) Recapitulation of Lerik1-3 × Fei-0 hybrid necrosis with genomic *RPP7b*^Lerik1-3^ construct in four-week-old T_1_ transgenic Fei-0 plants, grown at 23°C. *RPP7* genes were expressed under the control of the *RPP7b* promoter. Size bar corresponds to 1 cm. (C) Recapitulation of Lerik1-3 × Fei-0 hybrid phenotype by co-expression of *RPP7b*^Lerik1-3^ and *HR4*^Fei-0^ under control of their native promoters in Col-0 background. Three independent T_2_ transgenic lines grown at 23°C are shown. Size bar corresponds to 1 cm. (D) Distribution of different phenotypes in *A. thaliana* T_1_ plants with *RPP7b::RPP7b-FLAG* and *HR4::HR4*^Fei-0^*-HA* transgenes in different backgrounds, grown at 23°C. The numbers above indicate the total T_1_ plants scored according to the scale shown on the right. Size bar corresponds to 1 cm. (E) Four-week-old *A. thaliana* T_1_ transgenic plants with *RPP7b::RPP7b-FLAG* and *HR4::HR4*^Fei-0^*-HA* in different backgrounds, grown at 23°C. Size bar corresponds to 1 cm.

### Genetic requirements for HR4^Fei-0^/RPP7b-induced plant autoimmunity

Hybrid necrosis cases such as HR4^Fei-0^/RPP7b present an opportunity to investigate NLR signaling without the complication of effector-triggered modifications of NLR ligands. To determine whether HR4^Fei-0^/RPP7b provides an appropriate model for pathogen-dependent signaling by NLRs, we sought to determine whether the genetic interaction between RPP7b and HR4^Fei-0^ requires known downstream components of canonical NLR signaling. EDS1 and PAD4 typically act as positive regulators of TNL pathways, NDR1 mainly plays a positive role downstream of CNLs (Aarts et al., 1998; Wagner et al., 2013), and RAR1 and SGT1b control the stability of NLR proteins (Hubert et al., 2009; Zhang et al., 2010). Finally, NPR1 mediates the effects of the important signaling molecule salicylic acid (SA) (Cao et al., 1997; Ding et al., 2018). SA itself can be manipulated with mutations in the SA biosynthesis gene *SID2*, and to an even greater extent by expressing the bacterial enzyme NahG, which catabolizes SA (Nawrath and Métraux, 1999; Wildermuth et al., 2001). All of these components but NDR1 are essential for *RPW8.2*^Ms-0^ signaling during powdery mildew resistance (with *SID2* not having been tested) (Xiao et al., 2005), while *RPP7*^Col-0^ signaling during downy mildew resistance appears to rely on multiple, redundant pathways, such that *ndr1* mutation or a *NahG* transgene on their own have only weak effects on *RPP7*^Col-0^-mediated resistance (McDowell et al., 2000).

We transformed the *RPP7b::RPP7b-3*×*FLAG* / *HR4::HR4*^Fei-0^*-3*×*HA* construct into the aforementioned mutants. Necrosis was effectively suppressed not only by *eds1*, but also by *ndr1*, which sets autoimmunity triggered by *HR4*^Fei-0^/*RPP7b*^Lerik1-3^ apart from *RPW8.2*^Ms-0^- and *RPP7*^Col-0^-dependent resistance to powdery and downy mildew, respectively (Figure 1D). More similar to *RPW8.2*^Ms-0^ (Xiao et al., 2005), but different from what has been reported for *RPP7*^Col-0^ (McDowell et al., 2000), *NahG* strongly attenuated *HR4*^Fei-0^/*RPP7b*^Lerik1-3^-induced necrosis, while *npr1* and *sid2* did not (Figure 1E). The requirement for multiple disease resistance signaling components is consistent with HR4^Fei-0^/RPP7b hybrid necrosis mimicking conventional, pathogen-triggered signaling. At the same time, the differences between pathogen-triggered signaling initiated by *RPW8.2*^Ms-0^ and *RPP7*^Col-0^ as well as pathogen-independent signaling by *HR4*^Fei-0^/*RPP7b*^Lerik1-3^ suggest that not all RPW8/HR and RPP7 proteins act through the same pathways.

### RPP7b oligomerization induced by association with HR4^Fei-0^

Certain conventional NLRs work as heteromeric pairs, through interacting N-terminal domains (Césari et al., 2014; Tran et al., 2017; Williams et al., 2014). RPP7 proteins are canonical CNLs, while RPW8/HR proteins are non-NLR, CC_R_-only proteins. Co-immunoprecipitation (co-IP) indicated that HR4Fei-0 and RPP7b associate in *A. thaliana* cells (Figure 2A), and truncation analysis revealed that the CC and LRR domains of RPP7b are mainly responsible for interaction with HR4^Fei-0^ (Figure S2A). The CC and LRR domains of RPP7b can bind HR4^Fei-0^ directly, as demonstrated with in vitro pull-down assays (Figure 2B).

**Figure 2.**
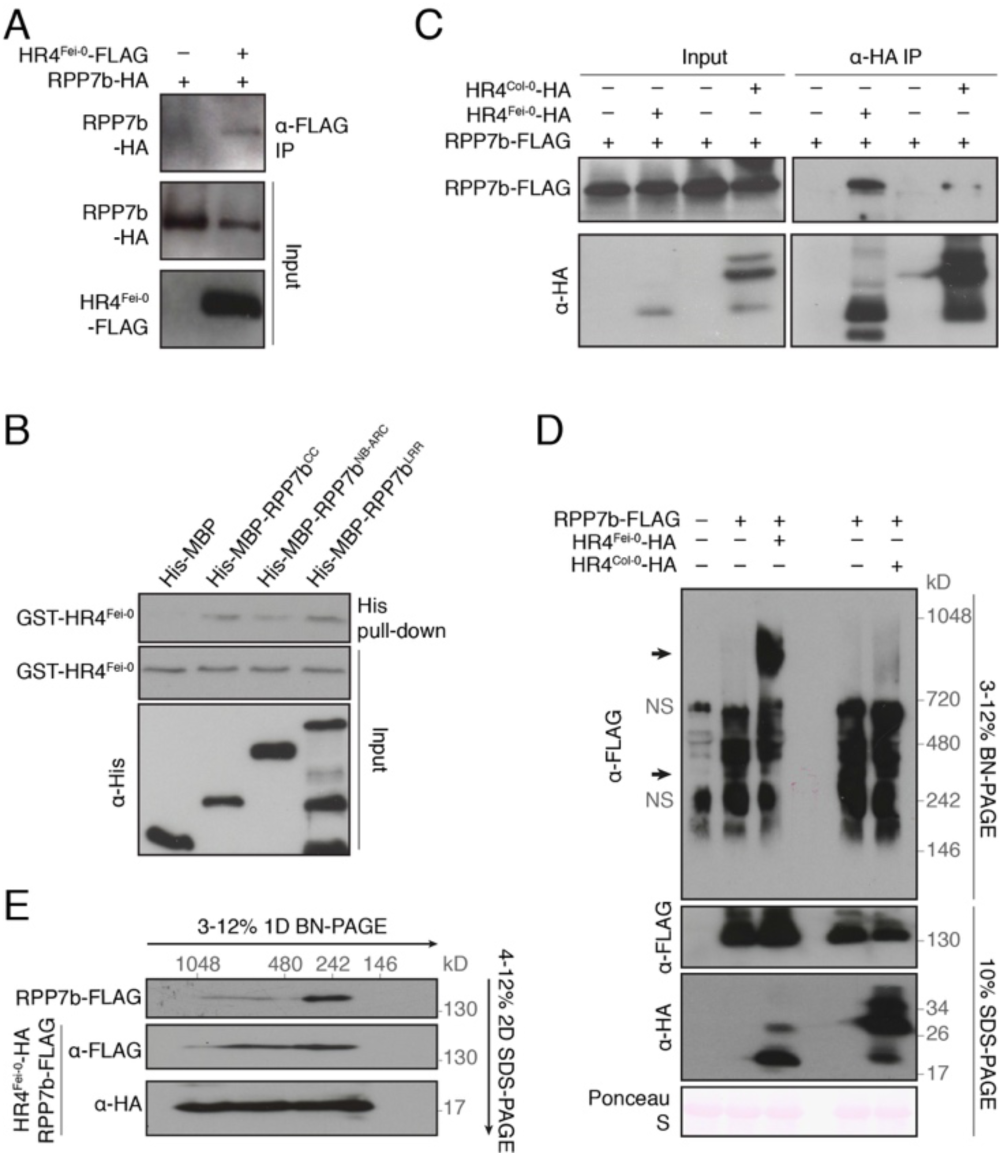
HR4^Fei-0^ associates with RPP7b and induces oligomerization of RPP7b. (A) HR4^Fei-0^ interacts with RPP7b in vivo, as shown by co-IP from *A. thaliana* protoplasts. (B) HR4^Fei-0^ interacts with RPP7b in vitro, as shown by pull-down assays with proteins purified from *E. coli*. (C) HR4^Fei-0^ interacts with RPP7b in vivo, as shown by co-IP from transgenic *A. thaliana* plants. Expression of RPP7b::RPP7b-FLAG and HR4^Fei-0^-HA or HR4^Col-0^-HA was induced by β-estradiol. (D) HR4^Fei-0^ induces oligomerization of RPP7b, as shown by BN-PAGE and SDS-PAGE. Arrows point to RPP7b complexes. NS indicates nonspecific bands. (E) Evidence for HR4^Fei-0^ and RPP7b residing in the same complex, as shown with second-dimension SDS-PAGE using lanes 2 and 3 of BN-PAGE in (D). Experiments were performed at least three times.

We further investigated the specificity of the physical interaction between RPP7b and HR4^Fei-0^ using *A. thaliana* plants carrying a *RPP7b-3×FLAG* transgene under control of its native promoter and a *HR4*^Fei-0^*-3*×*HA* or *HR4*^Col-0^*-3*×*HA* transgene under control of an estradiol-inducible promoter. Only the *HR4*^Fei-0^*-3*×*HA* but not the *HR4*^Col-0^*-3*×*HA* plants produced necrosis after estradiol treatment (Figure S2B). In co-IPs, we consistently observed that RPP7b associated with HR4^Fei-0^ more strongly than with HR4^Col-0^ (Figure 2C), consistent with HR4^Fei-0^/RPP7b interaction being a prerequisite for activation of immune signaling.

To investigate whether HR4^Fei-0^ and RPP7b form stable complexes in vivo, and to determine the size of such complexes, we turned to Blue Native polyacrylamide gel electrophoresis (BN-PAGE) of material extracted from transgenic *A. thaliana* plants. In the absence of HR4^Fei-0^, we detected an RPP7b complex of about 300 kD (Figure 2D) but not a RPP7b monomer, implying that RPP7b exists in the resting state in a homomeric complex (or in a heteromeric complex with another protein). RPP7b shifted to a larger complex of about 900 kD in the presence of HR4^Fei-0^ but not HR4^Col-0^ (Figure 2D). Analysis of complexes in a second dimension of SDS-PAGE further verified that HR4^Fei-0^ can promote higher-order oligomerization of RPP7b, with HR4^Fei-0^ being present in high-molecular-mass complexes (Figure 2E). Taken together, these results demonstrate that signaling by the plant NLR RPP7b requires formation of a higher-order oligomeric complex. This observation is consistent with cryo-EM studies of complexes of plant NLR ZAR1 and animal NLR NLRC4 (Hu et al., 2015; Wang et al., 2019b; Zhang et al., 2015).

### Requirement of P-loop function for HR4^Fei-0^-induced RPP7b oligomerization

An attractive model of plant NLR function has NLRs existing in an equilibrium between an active ATP-bound and an inactive ADP-bound state (Bernoux et al., 2016). The balance between the active and inactive state in turn is thought to be regulated by ligands that affect intramolecular interactions between different NLR domains (Ade et al., 2007; Bernoux et al., 2016; Moffett et al., 2002; Schreiber et al., 2016), consistent with the structures of inactive and active ZAR1 forms (Wang et al., 2019a, 2019b).

To investigate how the association with HR4^Fei-0^ might affect intra- and/or intermolecular interactions between different RPP7b domains, we carried out co-IP assays with material from *A. thaliana* protoplasts that co-expressed combinations of CC, NB-ARC or LRR domains of RPP7b. The LRR domain interacted strongly with CC and NB-ARC domains, and the NB-ARC domain interacted weakly with the CC domain (Figure 3A), indicating that all RPP7b domains contribute to intramolecular association. Furthermore, both the CC and LRR domains can self-associate (Figure 3A), suggestive of dimerization through homophilic intermolecular interactions.

**Figure 3.**
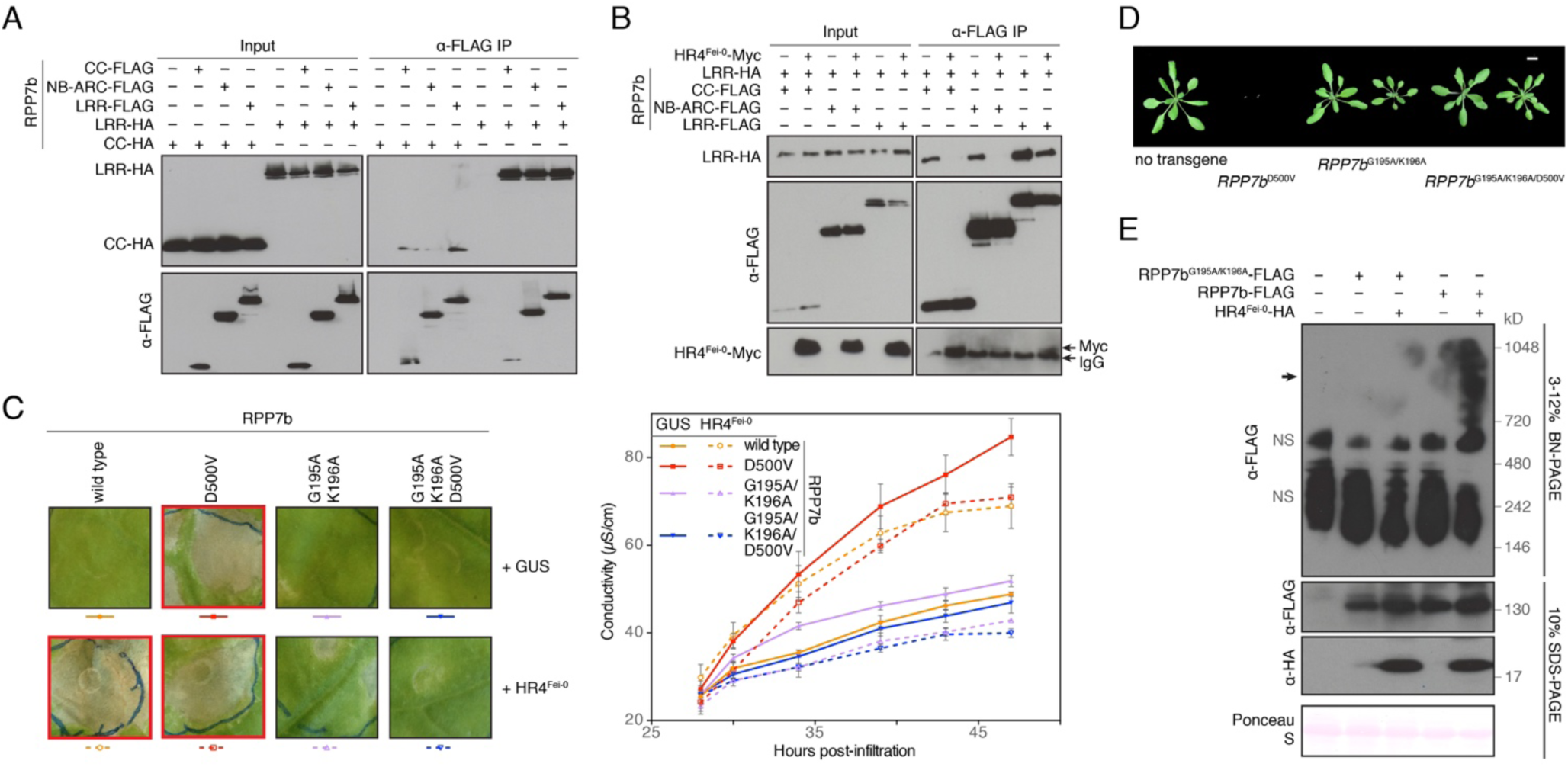
HR4^Fei-0^ induces oligomerization of RPP7b in a P-loop dependent manner. (A) Interaction between RPP7b domains, as shown by co-IP from *A. thaliana* protoplasts. (B) HR4^Fei-0^ weakens interactions between RPP7b domains, as shown by co-IP from *A. thaliana* protoplasts. (C) Analysis of RPP7b variants with P-loop and MHD motif mutations in *N. benthamiana*, seven days after *Agrobacterium* infiltration. *RPP7b* variants, all expressed under the control of the *RPP7b* promoter, were co-expressed with *HR4*^Fei-0^. Left, images of leaves; hypersensitive response is indicated by red frames. Experiments were performed at least three times. Right, ion leakage measurements of plants shown on left. Values are means ± SEM (n=3). Each sample contains eight leaf discs from different plants, and experiments were performed at least twice. (D) Analysis of P-loop and MHD motif mutants in *A. thaliana*. Four-week-old T_1_ transgenic plants with *RPP7b* variants expressed from *RPP7b* promoter in Fei-0 background, grown at 23°C. Size bar corresponds to 1 cm. (E) HR4^Fei-0^ induces oligomerization of RPP7b in a P-loop dependent manner, as shown by BN-PAGE and SDS-PAGE. Arrow points to RPP7b high-molecular weight complexes. NS indicates nonspecific bands.

Since HR4^Fei-0^ interacts directly with RPP7b and promotes formation of a higher-order complex, we next asked whether HR4^Fei-0^ affects intra- and/or intermolecular RPP7b interactions. We transiently expressed RPP7b domains in the absence or presence of HR4^Fei-0^ in *A. thaliana* protoplasts and performed co-IPs. We observed that HR4^Fei-0^ significantly weakened interactions between the LRR domain and either the CC or NB-ARC domain, whereas the homophilic interaction between LRR domains was unaffected by HR4^Fei-0^ (Figure 3B). In combination with the results from BN-PAGE, we conclude that HR4^Fei-0^ induces conformational changes of RPP7b that alter the interaction between its different domains, resulting in RPP7b oligomerization as a prerequisite for downstream immune signaling.

To determine whether the inferred conformational changes of RPP7b depend on an active ATP-binding P-loop motif (Sukarta et al., 2016; Tameling et al., 2006; Williams et al., 2011), we substituted the GMGGL<u>GK</u>T motif with GMGGL<u>AA</u>T. This mutation significantly attenuated RPP7b activity when co-expressed with HR4^Fei-0^ in *N. benthamiana* (Figure 3C). Conversely, changing the ADP-binding pocket motif MHD to MHV, known to render many NLRs constitutively active because it destabilizes the ADP-bound state (Sukarta et al., 2016; Tameling et al., 2006; Williams et al., 2011), conferred constitutive activity on RPP7b^D500V^ in the absence of the HR4^Fei-0^ partner (Figure 3C). However, it could not override the inactivating effects of the P-loop mutation, neither in the presence or absence of HR4^Fei-0^ (Figure 3C). We confirmed that this was not due to insufficient protein accumulation (Figure S3A), and we also confirmed the effects of the different mutations and their combination by transforming *RPP7b* mutant variants into the Fei-0 background (Figure 3D).

We further analyzed the interaction between P-loop mutated RPP7b^G195A/K196A^ and HR4^Fei-0^. The P-loop mutant RPP7b^G195A/K196A^ retained the ability to associate with HR4^Fei-0^, as shown by co-IP (Figure S3B). However, RPP7b^G195A/K196A^ did not form a higher-order complex in the presence of HR4^Fei-0^, as demonstrated with BN-PAGE (Figure 3E). We thus conclude that RPP7b oligomerization occurs in two steps, beginning with HR4^Fei-0^ binding to RPP7b, followed by P-loop dependent conformational change and finally formation of a higher-order complex.

NLR conformational changes expose functional domains for participation in downstream signaling (Wang et al., 2019a, 2019b), and CC domains alone can often trigger immune signaling when expressed on their own (Casey et al., 2016; Cesari et al., 2016; Maekawa et al., 2011b), even in the absence of an activating partner. To test whether the CC or any other RPP7b domain can induce a hypersensitive response, we transiently expressed CC, NB-ARC, LRR, CC-NB-ARC, and NB-ARC-LRR constructs in *N. benthamiana*. None of the domains of RPP7b elicited a hypersensitive response, regardless of the presence of HR4^Fei-0^ (Figure S3C and S3D). Furthermore, different from the full-length version of RPP7b, the autoactivating MHV mutation did not confer autoactivity on any of the truncated forms (Figure S3D). These results suggest that the integrity of all domains of RPP7b is required for functional complex assembly and initiation of signaling.

### Requirement of RPP7 LRRs for HR4^Fei-0^-induced RPP7 oligomerization

The LRR domains of plant NLRs are often responsible for pathogen recognition and correspondingly polymorphic. Comparison of RPP7b^Lerik1-3^ with other RPP7 variants (Van de Weyer et al., 2019) revealed various LRR domain insertions and deletions in the phylogenetic clade that includes RPP7b^Lerik1-3^ (Figure 4A). Co-expression of HR4^Fei-0^ with RPP7^Sf-2^, which has the same LRR domain as RPP7b^Lerik1-3^, elicited a hypersensitive response in *N. benthamiana*, albeit more weakly than co-expression of RPP7b^Lerik1-3^ (Figure 4B). Similarly, we found that RPP7^Mrk-0^, which is incompatible with RPW8.1^KZ10^, is distinguished from other RPP7 variants by a truncated LRR domain (Van de Weyer et al., 2019) (Figure S4A). The causality of the LRR truncation was confirmed with RPP7^Ty-1^, which is very similar to RPP7^Mrk-0^, but which does not have a truncated LRR domain and thus is not incompatible with RPW8.1KZ10 (Figure S4B-E). F_1_ plants derived from crosses between KZ10 and six other accessions having RPP7 homologs with truncated LRR domains developed strong hybrid necrosis (1001 Genomes Consortium, 2016) (Figure S4B). LRR truncation is, however, not sufficient for hybrid necrosis, as inferred from four further accessions carrying LRR-truncated RPP7 alleles that did not cause hybrid necrosis in crosses with KZ10 (Figure S4B).

**Figure 4.**
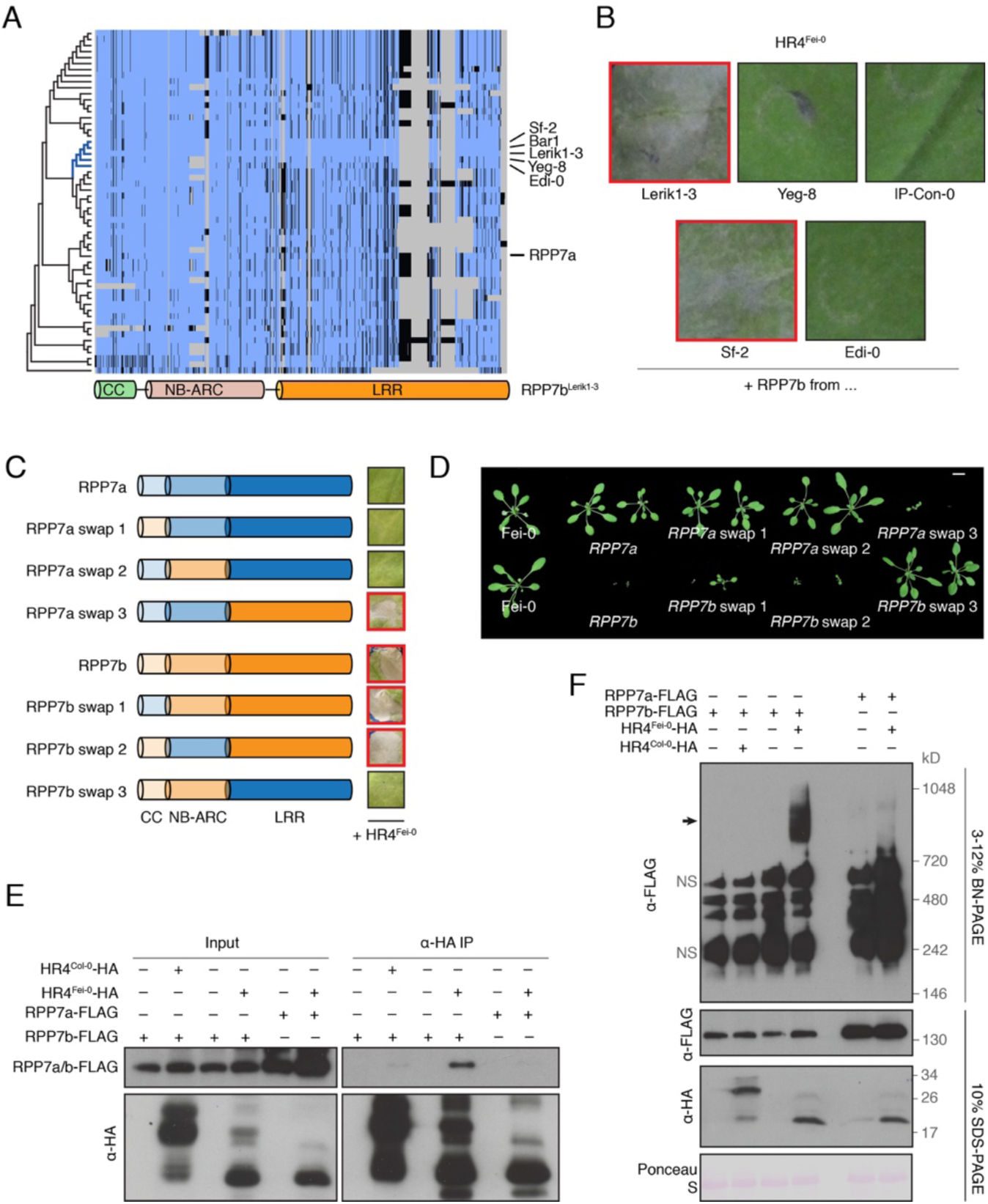
Requirement of RPP7 LRRs for HR4^Fei-0^-induced oligomerization of RPP7b. (A) Alignments of RPP7 related proteins from 64 accessions with RPP7b^Lerik1-3^ reveal sequence and structural polymorphisms, especially in the LRR domain. Blue, match; dark, mismatch; grey, gap. (B) *RPP7b*^Lerik1-3^-like alleles produce a hypersensitive response (indicated by red frames) when co-expressed with HR4^Fei-0^ in *N. benthamiana*, seven days after *Agrobacterium* infiltration. Experiments were performed at least three times. (C) Specific domain swaps between RPP7a and RPP7b, when co-expressed with HR4^Fei-0^, can induce hypersensitive response (indicated by red frames) in *N. benthamiana*, seven days after *Agrobacterium* infiltration. Experiments were performed at least twice. (D) Four-week-old T_1_ transgenic *A. thaliana* plants expressing chimeras from (C) in Fei-0 background, grown at 23°C. *RPP7* swaps were under the control of the *RPP7b* promoter. Size bar corresponds to 1 cm. (E) HR4^Fei-0^ specifically interacts with RPP7b in vivo, as shown by co-IP from transgenic *A. thaliana* plants. Expression of RPP7a-FLAG or RPP7b-FLAG under the control of its native promoter and HR4^Fei-0^-HA or HR4^Col-0^-HA was induced by β-estradiol. (F) HR4^Fei-0^ specifically induces the oligomerization of RPP7b, as shown by BN-PAGE and SDS-PAGE. Arrow points to RPP7b high-molecular weight complexes. NS indicates nonspecific bands. Experiments of (E) and (F) were performed at least twice.

To further dissect the function of different domains, we generated a series of RPP7a^Lerik1-3^/RPP7b^Lerik1-3^ chimeras. Swapping the CC or NB-ARC domains had no effect on either RPP7a or RPP7b activity (Figure 4C, S4F and S4G, swaps 1 and 2). In contrast, exchanging the LRR domains inactivated RPP7b (Figure 4C, RPP7b swap 3), while the RPP7b LRR domain was sufficient to impart necrosis-inducing activity on RPP7a (Figure 4C, RPP7a swaps 3). Transgenic *A. thaliana* plants further confirmed the results obtained with transient expression in *N. benthamiana* leaves (Figure 4D). Additional chimeras indicated that a largely intact LRR domain is required for RPW8/HR-dependent signaling and/or RPP7b protein stability (Figure S4F, S4H, S4I).

To investigate the mechanism underlying the specificity of HR4^Fei-0^/RPP7b signaling, we performed co-IPs with material from transgenic plants. These showed that HR4^Fei-0^ co-immunoprecipitated RPP7b more efficiently than RPP7a (Figure 4E), indicating that association of RPP7b with HR4^Fei-0^ is correlated with immune signaling. Preferential binding of HR4^Fei-0^ to the RPP7b LRR domain was further supported by co-IPs and pull-down assays (Figure S4J and S4K). Homology modeling of RPP7a and RPP7b LRR structures and superposition of these two structures suggested that the RPP7b LRR domain, which provides an interface for interaction with HR4^Fei-0^, has a more open conformation (Figure S4L). Finally, BN-PAGE showed that HR4^Fei-0^ cannot induce oligomerization of RPP7a (Figure 4F). We conclude that specific association between HR4^Fei-0^ and RPP7b, with the RPP7b LRR domain providing specificity for interaction with the HR4^Fei-0^ ligand, induces the oligomerization of RPP7b in vivo and initiates immune signaling.

### RPP7b-independent activity of HR4^Fei-0^

The differences in genetic requirements for RPW8/HR and RPP7 signaling on their own (McDowell et al., 2000; Xiao et al., 2005) prompted us to test whether HR4^Fei-0^ proteins might have activity independently of RPP7 partners. Indeed, we found that while moderate expression of *HR4*^Fei-0^ in *N. benthamiana* could not induce a hypersensitive response without its *RPP7b* partner (Figure 1A), strong overexpression of *HR4*^Fei-0^ could (Figure 5A). Further evidence for RPP7b-dependent and -independent activities of HR4^Fei-0^ came from structure-function studies: While even small N-terminal deletions compromised the stand-alone activity of HR4^Fei-0^, these did not affect or even increased activity of HR4^Fei-0^ in the presence of RPP7b (Figure 5A). In contrast, C-terminal deletions had similar effects in the presence or absence of RPP7b (Figure 5A).

**Figure 5.**
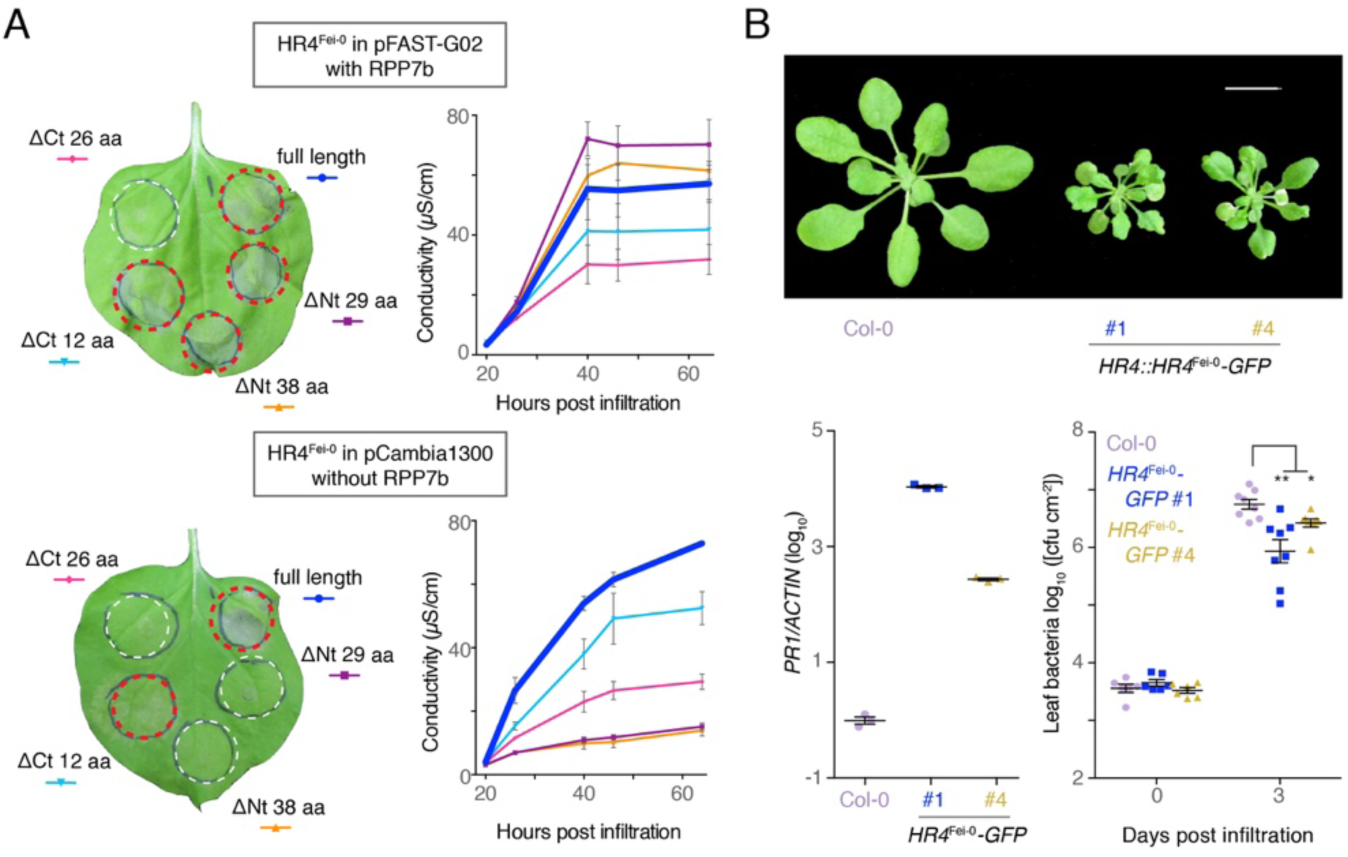
RPP7b-independent activity of HR4^Fei-0^. (A) Left, hypersensitive response (red circles) induced by expression of full-length or truncated HR4^Fei-0^ variants using pFAST-G02 for moderate expression (top) and pCambia1300 for strong overexpression (bottom), and with or without RPP7b in *N. benthamiana*, four days after *Agrobacterium* infiltration. For control for pFAST-G02 expression of HR4^Fei-0^ in the absence of RPP7b, see Figure 1A (experiment 7). Right panels, ion leakage measurements of plants shown on left. Values are means ± SEM (n=3). Each sample contains six leaf discs from different plants. Experiments were performed at least twice. (B) *HR4::HR4*^Fei-0^*-GFP* reduces plant growth, induces *PR1* defense marker and suppresses the bacterial model pathogen *Pseudomonas syringae* pv. *tomato* DC3000 (*P < 0.05, **P < 0.01, Student’s *t* test; n = 8).

To further investigate RPP7b-independent activity of HR4^Fei-0^, we transformed a fusion of HR4^Fei-0^ with GFP, a large tag that can promote dimerization (Shaner et al., 2005), into the Col-0 background. We found that *HR4::HR4*^Fei-0^*-GFP* plants were often smaller, a phenotype shared with other autoimmune mutants (Figure 5B). These plants also had increased expression of the defense marker *PR1*, and proliferation of the model pathogen *Pseudomonas syringae* DC3000 was substantially reduced (Figure 5B). Together, these observations suggest that HR4^Fei-0^ can trigger (auto)immunity in an RPP7b-independent manner.

### Aggregation of HR4^Fei-0^ contributes to its cell death activity

While RPP7b is not required for cell death induced by HR4^Fei-0^ overexpression or enhanced HR4^Fei-0^ dimerization, we could not exclude that HR4^Fei-0^ in such contexts associates with other, unknown NLRs. To directly test the ability of HR4^Fei-0^ to compromise cell growth without any NLRs, we therefore turned to a heterologous system, expression in *E. coli*, which has also been used to investigate the cytotoxic activity of the fungal HELL domain protein Het-S (Seuring et al., 2012). Expression of HR4^Fei-0^ strongly suppressed bacterial growth, and eventually reduced bacterial number, while HR4^Col-0^ had more modest effects on bacterial growth (Figure 6A). These experiments confirmed cytotoxicity of HR4^Fei-0^, and to a lesser extent of HR4^Col-0^, in a heterologous system.

**Figure 6.**
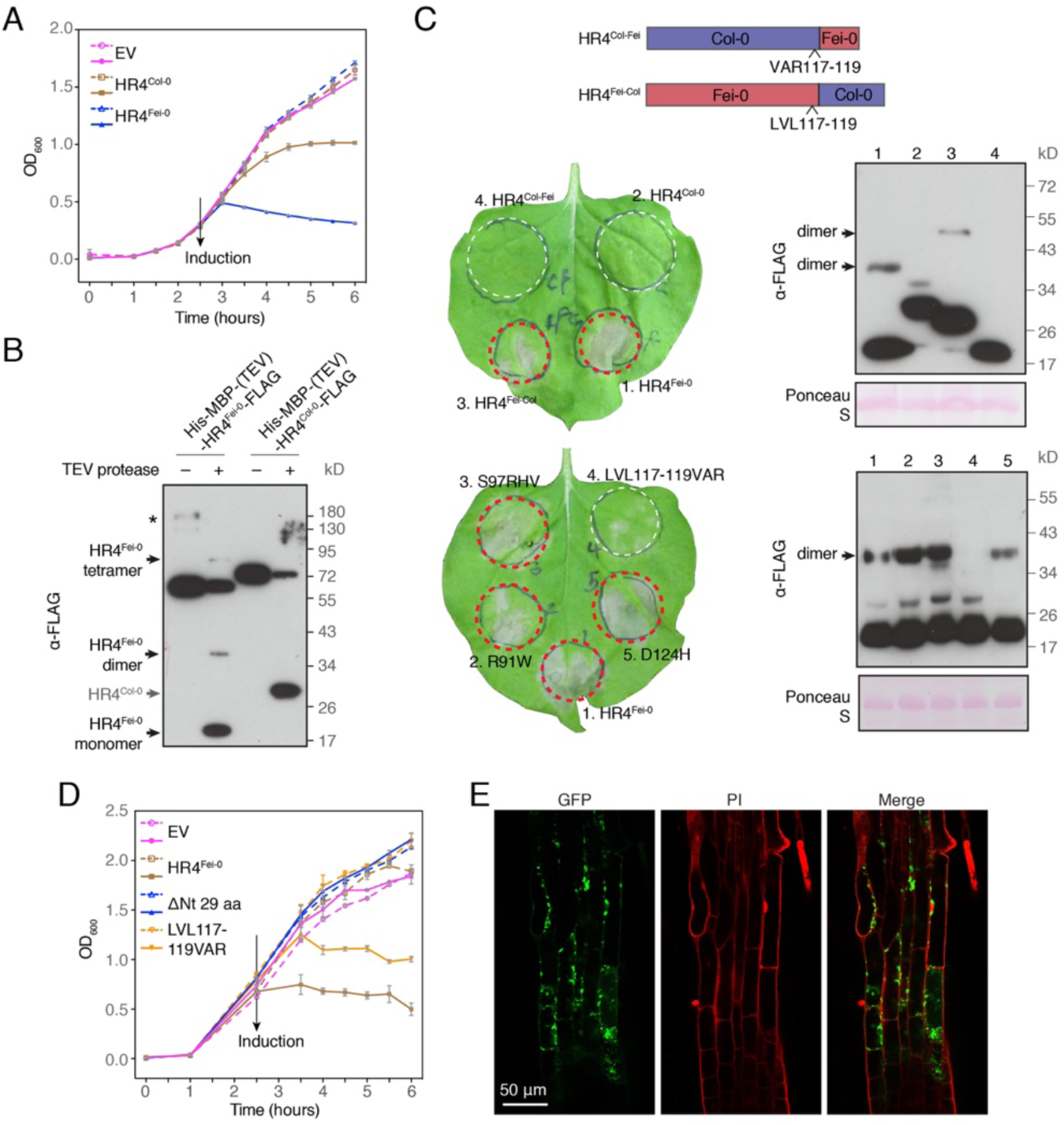
Aggregation of HR4^Fei-0^ contributes to its cytotoxic activity. (A) Expression of HR4^Fei-0^ is toxic to *E. coli*. Bacterial growth was determined by OD_600_ measurement; expression of heterologous proteins was induced (solid lines) with 0.2 mM IPTG at 37°C at the same time for all cultures. Values are means ± SEM (n=3). EV, empty vector. (B) Recombinant HR4^Fei-0^ but not HR4^Col-0^ protein forms SDS-resistant dimers and tetramers. The same amount, 3 μg, of His-MBP-(TEV) fusion proteins was incubated without or with 0.1 μg TEV protease overnight at 4°C. Compare major bands at 60-70 kD for assessment of TEV protease cleavage efficiency. Asterisk indicates fusion protein oligomers. (C) HR4 domain swaps and targeted mutations. Left, hypersensitive response (red circles) induced by overexpression of domain swaps between HR4^Fei-0^ and HR4^Col-0^ (top) or mutated HR4^Fei-0^ (bottom) in *N. benthamiana*, four days after *Agrobacterium* infiltration. Right, protein accumulation for experiments shown of left. SDS-resistant dimers are indicated. Only the LVL117-119VAR mutation abrogates hypersensitive response and dimer formation. (D) Mutations of HR4^Fei-0^ that inhibit its cytotoxic activity in *E. coli*. Bacterial growth was determined by OD_600_ measurement; expression of heterologous proteins was induced (solid lines) with 0.2 mM IPTG at 37°C at the same time for all cultures. Values are means ± SEM (n=3). EV, empty vector. (E) Confocal images of an *A. thaliana* transgenic line expressing inducible HR4^Fei-0^-GFP after 10 h treatment with β-estradiol. The plasma membrane was stained with propidium iodide (PI).

Het-S can form amyloidal aggregates that break down into detergent-resistant dimers and/or trimers (Mathur et al., 2012). To test whether HR4 behaved similarly, we purified His-MBP-HR4 fusion proteins from *E. coli* and analyzed the proteins after removal of the MBP tag. We found that HR4^Fei-0^ could form stable, SDS-resistant dimers or tetramers, but HR4^Col-0^ could not (Figure 6B). In addition, while both fusion proteins contained the same TEV protease cleavage site, the His-MBP-HR4^Fei-0^ fusion, was more resistant to TEV protease digestion than His-MBP-HR4^Col-0^. Finally, SDS-resistant dimers of HR4^Fei-0^ but not of HR4^Col-0^ were observed in *N. benthamiana*, paralleling their ability to induce cell death (Figure 6C).

In contrast to the critical role of HR4^Fei-0^ C-terminal repeats in activation of hybrid necrosis in combination with RPP7b (Barragan et al., 2019), we found that replacing the C-terminal repeats of HR4^Fei-0^ with the HR4^Col-0^ repeats (HR4^Fei-Col^) only weakly affected its ability to form SDS-resistant dimers and to elicit cell death on its own. In contrast, swapping the N-terminal domain (HR4^Col-Fei^) greatly reduced formation of SDS-resistant dimers and abrogated the ability to induce cell death (Figure 6C, top, and S6B). These results suggest that the N-terminal domain of HR4^Fei-0^ determines its dimerization ability, and that dimerization in turn is required for its cytotoxic activity. Aggregation only, however, does not appear to be sufficient for the induction of cell death, since small N-terminal and C-terminal truncations did not impair the formation of SDS-resistant dimers (Figure S6A), but did affect the cytotoxic activity of HR4^Fei-0^ in planta (Figure 5A). We performed more fine-scale swaps and found that exchange of three consecutive amino acids, L117V, V118A, L119R, in the HR4^Fei-0^ background reduced hypersensitive response and dimer formation in *N. benthamiana* (Figure 6C, bottom). In agreement, cytotoxic activity of HR4^Fei-0^ in *E. coli* was also attenuated by the LVL117-119VAR mutation (Figure 6D). The N-terminal 29 amino acid deletion, which eliminated hypersensitive response in *N. benthamiana* (Figure 5A, bottom), also eliminated cytotoxic activity in *E. coli* (Figure 6D). Together, these results indicate that cell death-inducing activity of HR4^Fei-0^ in planta and in bacteria is structure dependent. The LVL117-119 residues are located near the border of the N-terminal domain and the C-terminal repeats (Figure S6B), reminiscent of the short overlap between the HeLo and PFD domains of Het-S (Greenwald et al., 2010).

Finally, using a transgenic *A. thaliana* line that expresses the dimerization prone HR4^Fei-0^-GFP fusion protein under the control of an inducible promoter, we observed that HR4^Fei-0^-GFP accumulated as various-sized puncta that were localized at or close to the plasma membrane (Figure 6E), suggestive of aggregates formed in planta. In contrast, we could not detect such puncta with control transgenic lines expressing GFP only or HR4^Col-0^-GFP (Figure S6C). Some diffuse HR4^Fei-0^-GFP fluorescence was observed in the cytoplasm and around the nucleus (Figure S6C), which may correspond to Z-membranes, an artificial organelle formed by oligomerization of integral membrane proteins (Gong et al., 1996).

## DISCUSSION

We have identified a two-pronged immunity signaling module with the enigmatic RPW8/HR proteins at its core. We took advantage of genetics and biochemistry to reveal the activation mechanism underlying HR4^Fei-0^/RPP7b-induced plant immunity. Our data support a model in which HR4^Fei-0^ acts as a ligand that disrupts intramolecular association of RPP7b, which leads to its unfolding, thereby initiating oligomerization and finally immune signaling by RPP7b. In addition, we discovered that HR4^Fei-0^ on its own can trigger plant immunity, with distinct portions of the HR4^Fei-0^ molecule being involved in RPP7b-dependent and -independent immune signaling.

The first RPW8/HR protein to be functionally characterized was RPW8.2, because it provides immunity to powdery mildew fungi (Xiao et al., 2001). Constitutive expression of other RPW8/HR proteins endows plants with broad-spectrum resistance to both fungi and downy mildew oomycetes (Ma et al., 2014). Such broad-spectrum resistance, along with localization of several of these proteins to extrahaustorial membranes that form at the interface of plant epidermal cells and invading fungal cells, is suggestive of RPW8/HR proteins providing primary defenses against filamentous pathogens. However, the exact mode of action of different RPW8/HR proteins may differ, as indicated by RPW8.1 accumulating around chloroplasts of mesophyll cells next to epidermal cells contacted by the fungus (Berkey et al., 2017; Wang et al., 2009),

An important observation is that HR4^Fei-0^ can cause autoimmunity even in the absence of the RPP7b partner. Homology modelling of RPW8/HR proteins has revealed striking similarity with MLKL and HELL domain proteins, executors of cell death in animals and fungi (Bentham et al., 2018; Daskalov et al., 2016; Greenwald et al., 2010; Hildebrand et al., 2014; Murphy et al., 2013; Seuring et al., 2012; Su et al., 2014). The activity of both MLKL and Het-S is regulated by sequences immediately C-terminal to the domain shared by these proteins, corresponding to the CC_R_ domain of RPW8/HR proteins (Chen et al., 2014; Hildebrand et al., 2014; Murphy et al., 2013; Seuring et al., 2012; Su et al., 2014; Wang et al., 2014). Similar to the HELL domain protein Het-S (Seuring et al., 2012), expression of HR4^Fei-0^ is cytotoxic in bacteria, with RPP7b-independent activity of HR4^Fei-0^ likely being due to protein aggregation that is nucleated by C-terminal repeats (Figure 6). Variation in C-terminal repeats is also causal for hybrid necrosis activity of RPW8/HR variants in combination with specific RPP7 partners (Barragan et al., 2019), strongly suggesting that these repeats affect RPW8/HR conformation. While it was previously not possible to detect HR4 from Col-0 as a YFP fusion protein, even when expressed from a strong constitutive promoter (Berkey et al., 2017), we have been able to detect punctate aggregates that localize to the plasma membrane when HR4^Fei-0^ was expressed from an inducible promoter. Given that other RPW8/HR proteins localize to membranes, we propose that at least HR4^Fei-0^ and likely other proteins in this family have a direct role in triggering cell death by disrupting membrane integrity. This in turn may activate danger signals, including damage-associated molecular patterns (DAMPs), which would explain why RPW8/HR activity relies at least in part on well known immunity signaling pathways (Xiao et al., 2005).

Given that our work points to a direct role of HR4^Fei-0^ in inducing cell death (Figure 5 and 6), an attractive hypothesis is that because RPW8/HR proteins provide primary defenses against pathogens, they are targets of yet to be identified pathogen effectors and as such guardees that are monitored by NLRs including RPP7 as guards (Jones et al., 2016). We propose that RPW8/HR hybrid necrosis proteins such as RPW8.1^KZ10^ and HR4^Fei-0^ mimic other RPW8/HR proteins that have been modified by effectors and that are erroneously recognized by matching RPP7 variants. That the genetic requirements for *RPP7*^Col-0^ signaling in *Hpa* resistance (McDowell et al., 2000) differ from the genetic requirements for *HR4*^Fei-0^/*RPP7b*^Lerik1-3^ autoimmunity (Figure 1D) suggests, however, that not all *RPP7* homologs depend on *RPW8*/*HR* partners. This is also consistent with the observation that *RPP7*^Col-0^-mediated resistance to *Hpa* race Hiks1 does not require *HR4*^Col-0^ (Barragan et al., 2019). There is precedence for closely related NLR proteins or even the same NLR protein associating with different ligands or responding to different effectors (Lewis et al., 2013; Seto et al., 2017; Wang et al., 2015).

Self-association plays an important role in the activation of plant NLR proteins (Bernoux et al., 2011; Casey et al., 2016; Cesari et al., 2016; Maekawa et al., 2011b; Zhang et al., 2017b). The recent cryo-EM structures of plant ZAR1 and animal NLRC4 have greatly advanced our understanding of the biochemical mechanisms underlying NLR signaling (Hu et al., 2015; Wang et al., 2019a, 2019b; Zhang et al., 2015), even though the ZAR1 resistosome (or any other resistosome) has not yet been observed in vivo. In the in vitro ZAR1 resistosome, ZAR1 and RKS1 exist as a heterodimer in the resting, inactive state. Similarly, in the NLRC4 inflammasome complex, both NLRC4 and the sensor NAIPs exist as monomers in the resting, inactive state. Upon activation, these plant and animal NLRs form higher-order complexes: in the NLRC4 case containing one NAIP and ten NLRC4 molecules, in the ZAR1 case containing five ZAR1 molecules. Our BN-PAGE results indicate that binding of RPP7b to its autoactive ligand HR4^Fei-0^ induces oligomerization and formation of a higher-order complex, likely with predominantly six to seven RPP7b-HR4^Fei-0^ subunits (∼900 kD) (Figure 3D and 5F). We could not detect RPP7b in its monomeric form in vivo, regardless of the presence of HR4^Fei-0^. The size of inactive RPP7b protein as inferred by BN-PAGE is approximately twice its monomeric size (Figure 2D), suggesting that, similar to other plant NLRs, RPP7b hetero- or homodimers are formed before activation. In plant and animal NLR hetero-pairs, P-loop function of the sensor NLRs, such as RGA5, RRS1, and NAIP5, is dispensable for their activity (Césari et al., 2014; Williams et al., 2014; Zhao et al., 2011). In the ZAR1 resistosome, modification of the ZAR1-RKS1 complex by the PBL2^UMP^ ligand triggers a large conformational change, resulting in allosterically driven release of ADP bound to the NB-ARC domain, and subsequent assembly of a pentameric ZAR1-RKS1-PBL2^UMP^ oligomer. In analogy with the activation of the ZAR1 resistosome, we propose that HR4^Fei-0^ induces RPP7b to form a higher-order complex in a P-loop dependent manner through direct interaction between the RPP7b LRR domain and HR4^Fei-0^. For this to occur, matching partners on both the RPW8/HR and on the RPP7 side are required.

In summary, we have investigated interactions between two arms of the plant immune system that have not been directly linked before, the CNL RPP7 and the non-NLR RPW8/HR proteins. We demonstrate that HR4^Fei-0^ protein can affect immune signaling both on its own and through its RPP7b partner. Our findings provide an excellent example of the usefulness of autoimmunity as a platform for mechanistic investigation of immune signaling.

## METHODS

### Plant material and growth conditions

Seeds of *A. thaliana* accessions and *N. benthamiana* were from laboratory stocks. *A. thaliana* plants were grown on soil at 23°C or 16°C, at 65% relative humidity under 16/8 h (long days) or 8/16 h (short days) light/dark photoperiods with light (125 to 175 μmol m^-2^ s^-1^) provided by a 1:1 mixture of Cool White and Gro-Lux Wide Spectrum fluorescent lights (Luxline plus F36W/840, Sylvania, Erlangen, Germany). *N. benthamiana* plants were grown in short days at 23°C with same light intensity and humidity.

### Constructs and transgenic plants

To generate constructs for protoplast transformation, fragments from cDNA were amplified by PCR and inserted into the pUC19-35S-FLAG-RBS and pUC19-35S-HA-RBS vectors (Li et al., 2005).

To generate constructs for recombinant proteins, HR4^Fei-0^ (30-Ct aa) or full length HR4^Fei-0^ was amplified and cloned into the pGEX-6p-1 or pET-28a vector, respectively. RPP7a/b^CC^ (1-146 aa), RPP7a/b^NB-ARC^ (147-520 aa), RPP7a/b^LRR^ (521-Ct aa), were amplified from cDNA and inserted into the pETM-41 vector.

To generate genome editing vectors, plasmids with an *A. thaliana* codon optimized *Cas9* were assembled using the GreenGate system (Wu et al., 2018a); *Cas9*-free transgenic seeds were selected by mCherry fluorescence in their seed coats.

To generate RPP7 and RPW8 constructs with plant promoters for transient expression or for stable transformation, all *RPP7* variants with the *RPP7b* promoter (1,491 bp), *RPW8.1*^KZ10^ with native promoter (1,122 bp), and *HR4*^Fei-0^ with native promoter (1,178 bp) were amplified from corresponding genomic DNA. Fragments were inserted into a modified Gateway TOPO entry vector by ligation or Gibson assembly (New England Biolabs, NEB), and then transferred into pGWB or pFAST vectors (Nakagawa et al., 2007; Shimada et al., 2010).

To generate constructs containing both RPP7 and HR4, fragments of genomic RPP7 with a promoter sequence from RPP7b (1,491 bp) were inserted into the Gateway TOPO entry vector. The HR4 fragments with either a native promoter (1,178 bp) or XVE inducible module from pER8 (Zuo et al., 2000) were then inserted into the above vector. Finally, the combined sequences were moved into the pFAST-G01 vector (Shimada et al., 2010).

Point mutations were introduced into the entry constructs described above by site-directed mutagenesis. Chimeric constructs were generated by Gibson assembly (New England Biolabs, NEB) based on above mentioned entry vectors.

### Transient expression and conductivity assay

Four-week-old *N. benthamiana* plants were used for *Agrobacterium tumefaciens*-mediated transient expression. *A. tumefaciens* with different T-DNA vectors were grown overnight at 28°C to OD_600_ around 1.5. Bacterial cells were harvested and resuspended in the induction buffer (10 mM MES [pH 5.7], 10 mM MgCl_2_ and 150 μM acetosyringone) with OD_600_ of 0.5, and incubated in induction buffer at 28°C in a shaker for 2 h. Bacterial inocula were mixed at 1:1 vol/vol ratio and infiltrated using a needleless syringe.

For conductivity assays, eight independent *N. benthamiana* leaves were injected with different bacteria. Three discs from each plant were collected 20 hours post-infiltration (hpi) and floated on 50 mL ddH_2_O for 30 min. Leaf discs were transferred into tubes with 5 mL ddH_2_O. Conductivity was measured using an Orion Conductivity Meter (Thermo Scientific, Beverly, MA, USA).

### Co-immunoprecipitation assays

For anti-FLAG co-IPs with material from *A. thaliana* protoplasts, isolation and transformation of protoplasts were performed as previously described (Yoo et al., 2007). Total protein was extracted with 600 μL extraction buffer (50 mM HEPES [pH 7.5], 150 mM KCl, 1 mM EDTA [pH 8.0], 0.4% Triton X-100, 1 mM DTT, proteinase inhibitor cocktail), and incubated with 30 μL ANTI-FLAG M2 Affinity agarose (Sigma-Aldrich, MO, USA) for 4 h at 4°C. Agarose was washed seven times with extraction buffer without proteinase inhibitor cocktail, and bound protein was eluted with 60 μL of 0.5 mg/mL 3×FLAG peptide (Sigma-Aldrich, MO, USA) for 1 h at 4°C. Proteins were separated by 10% SDS-PAGE and detected by immunoblot.

For anti-HA co-IP assays with material from *A. thaliana* plants, 100 mg of 10-day-old seedlings treated with or without 10 μM β-estradiol for 12 h were ground. Total protein was extracted with 600 μL TBS extraction buffer (50 mM Tris-HCl [pH 7.5], 150 mM NaCl, 1 mM EDTA [pH 8.0], 0.4% Triton X-100, proteinase inhibitor cocktail), and incubated with 25 μL anti-HA magnetic beads (Pierce, MA, USA) for 2 h at 4°C. Beads were washed seven times with TBS extraction buffer without proteinase inhibitor, and the bound protein was eluted with 60 μL of 0.1 M glycine (pH 2.2) for 10 min at room temperature and neutralized with 9 μL 1 M Tris-HCl (pH 8.3). Proteins were separated by 10% SDS-PAGE and detected by immunoblot.

### Blue Native-PAGE

Blue native polyacrylamide gel electrophoresis (BN-PAGE) was performed using the Bis-Tris NativePAGE system from Invitrogen (Carlsbad, CA, USA) according to the manufacturer’s instructions. Briefly, eight 10-day-old seedlings were collected and ground in 1×NativePAGE Sample Buffer (Invitrogen) containing 1% n-dodecyl β-D-maltoside (DDM) and protease inhibitor cocktail, followed by 13,000 rpm centrifugation for 20 min at 4°C. 20 μL supernatant mixed with 1 μL 5% G-250 Sample Additive was loaded and run on a NativePAGE 3-12% Bis-Tris gel. Native gels were transferred to PVDF membranes (Millipore, Darmstadt, Germany) using NuPAGE Transfer Buffer, followed by protein blotting. For the second dimension of electrophoresis, a 5.7 cm strip of BN-PAGE gel was incubated in Laemmli sample buffer (50 mM Tris-HCl [pH 6.8], 100 mM DTT, 2% (w/v) SDS, 0.1% bromophenol blue, 10% (v/v) glycerol) for 10 min, microwaved for 20 sec, and then rotated for another 5 min before loading the strip into the well of a NuPAGE 4-12% Bis-Tris protein gel (Invitrogen).

### Recombinant protein purification and in vitro His pull-down

Recombinant proteins were expressed in *E. coli* C41 cells. Bacterial cultures were induced by 0.4 mM IPTG at OD_600_ of 0.4 at 20°C for 16 h. GST-tagged and His-tagged proteins were affinity-purified using glutathione agarose beads (GE Healthcare, Pittsburgh, PA, USA) and Ni-NTA affinity agarose beads (QIAGEN, Venla, Netherlands) respectively, according to the manufacturer’s instructions. Finally, the buffer of recombinant proteins were changed into a buffer containing 25 mM Tris-HCl (pH 7.5), 100 mM NaCl, and 1 mM DTT using Amicon Ultra-15 Centrifugal Filter Units (Millipore, Darmstadt, Germany).

For His pull-downs, 0.5 µg His-MBP-tagged proteins, 2 µg GST-HR4^Fei-0^, 20 µL Ni-NTA affinity agarose beads were mixed in 500 µL buffer (25 mM Tris-HCl [pH 7.5], 150 mM NaCl, 0.2% Triton X-100, 20 mM imidazole) and incubated at 4°C on a rotator for 1 h. Agarose beads were washed seven times with the same buffer, and bound proteins were eluted by 60 µL elution buffer (25 mM Tris-HCl [pH 7.5], 150 mM NaCl, 250 mM imidazole) for 15 min at 4°C. Proteins were detected by immunoblot using anti-GST (Santa Cruz Biotechnology, Santa Cruz, CA, USA) and anti-His (Sigma-Aldrich, St. Louis, MO, USA) antibodies.

### Phylogenetic analysis

RPP7 protein sequences were identified from an NLR Ren-seq data set (Van de Weyer et al., 2019) (http://ann-nblrrome.tuebingen.mpg.de/annotator/index) and aligned with Clustalx-2.1, either as full-length protein or trimmed NB-ARC domain. MEGA6 was used to reconstruct phylogenetic trees using Neighbor-Joining (NJ) method. Node confidence in NB-ARC domain-based tree was assessed by bootstrapping with 1,000 replicates.

### Confocal microscopy

Transgenic 7-day-old *A. thaliana* seedlings were treated with β-estradiol for 10 h and imaged with a Zeiss LSM780NLO confocal microscope. Fluorescence was excited with the following wavelengths: 405 nm for 4’,6-diamidin-2-phenylindol (DAPI), 488 nm for GFP, and 561 nm for propidium iodide (PI). Fluorescent signals were captured within the following emission wavelengths: 420 to 480 nm for DAPI, 500 to 540 nm for GFP, and 580 to 650 nm for PI. For DAPI and PI staining, seedings were submerged in 10 mg/L PI and 1 mg/L DAPI for 5 min, and then washed briefly in water before imaging.

### Accession numbers

Sequences of *RPP7*^Lerik1-3^, *RPP7a*^Lerik1-3^, *RPP7b*^Lerik1-3^ and *RPP7*^Mrk-0^ have been deposited in GenBank under accession numbers MK776953-MK776956.

## ACKNOWLEDGMENTS

We are most grateful to Cristina Barragan, Eunyoung Chae, Faird El-Kasmi, Thomas Lahaye, Thorsten Nürnberger, Rebecca Schwab, Bridgit Waithaka and Wangsheng Zhu for discussion and critical reading of the manuscript, to Cristina Barragan, Eunyoung Chae and Rui Wu for sharing plasmids and sequence information, and Johannes Stuttmann for sharing *eds1-12* seeds. Supported by an EMBO Long-term Fellowship (969-2016 to L.L), an HFSP Long-term Fellowship (LT000314/2017-L to L.L), ERC Advanced Grant IMMUNEMESIS (340602), the DFG through Collaborative Research Center CRC1101 and the Max Planck Society.

## AUTHOR CONTRIBUTIONS

LL and DW designed the study. LL, AH and KW performed the experiments. LL and DW analyzed the results. LL and DW wrote the paper.

## DECLARATION OF INTERESTS

The authors declare no competing interests.

## Supplemental Information

**Figure S1.**
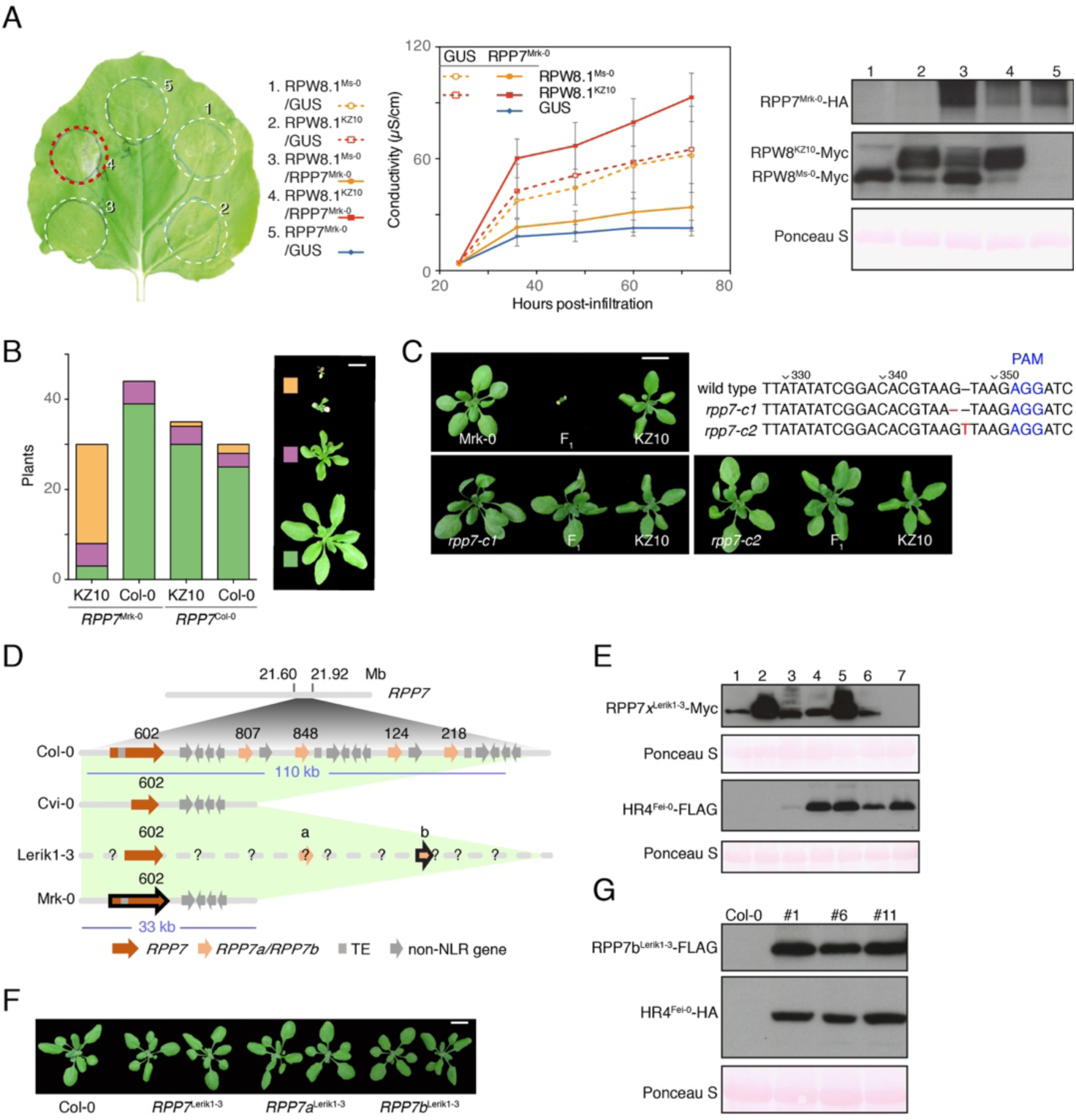
Identification of two distinct *RPP7* alleles as causal for hybrid necrosis. (A) Left, hypersensitive response (indicated by red frames) induced by co-expression of RPP7^Mrk-0^ with RPW8^KZ10^ in *N. benthamiana*, four days after *Agrobacterium* infiltration. Experiments were performed at least three times. Middle, Ion leakage measurements of plants shown on left. Values are means ± SEM (n=3). Each sample contains eight leaf discs from different plants, and experiments were performed at least twice. Right, protein accumulation as determined by immunoblot for experiments on left. (B) Recapitulation of Mrk-0 × KZ10 hybrid phenotype in four-week-old *A. thaliana* T_1_ transgenic plants, grown in 23°C. Size bar corresponds to 1 cm. (C) Requirement of *RPP7*^Mrk-0^ for hybrid necrosis demonstrated with CRISPR/Cas9-induced mutants, grown at 16°C. Two of the KZ10 control plants shown are identical, because F_1_s were grown at the same time. Size bar corresponds to 1 cm. Mutations generated by CRISPR/Cas9 methodology are indicated. (D) *RPP7* clusters of four *A. thaliana* accessions. The two causal alleles in Lerik1-3 and Mrk-0 are outlined in black. Numbers above arrows indicate At1g58**602**, At1g58**807**, At1g58**848**, At1g59**124** and At1g59**218**. The DNA sequences of At1g58807^Col-0^ and At1g59124^Col-0^ are identical, as are the DNA sequences of At1g58848^Col-0^ and At1g59218^Col-0^. TE, transposon. (E) Protein accumulation as determined by immunoblot for experiments in Figure 1A; see Figure 1A for numbering. (F) Four-week-old T_1_ transgenic plants expressing *RPP7* genes from Lerik1-3 in the neutral Col-0 background, grown at 23°C. (G) Protein expression for three transgenic lines shown in Figure 1C with *RPP7b::RPP7b*^Lerik1-3^*-FLAG* and *HR4::HR4*^Fei-0^*-HA* in Col-0 background.

**Figure S2.**
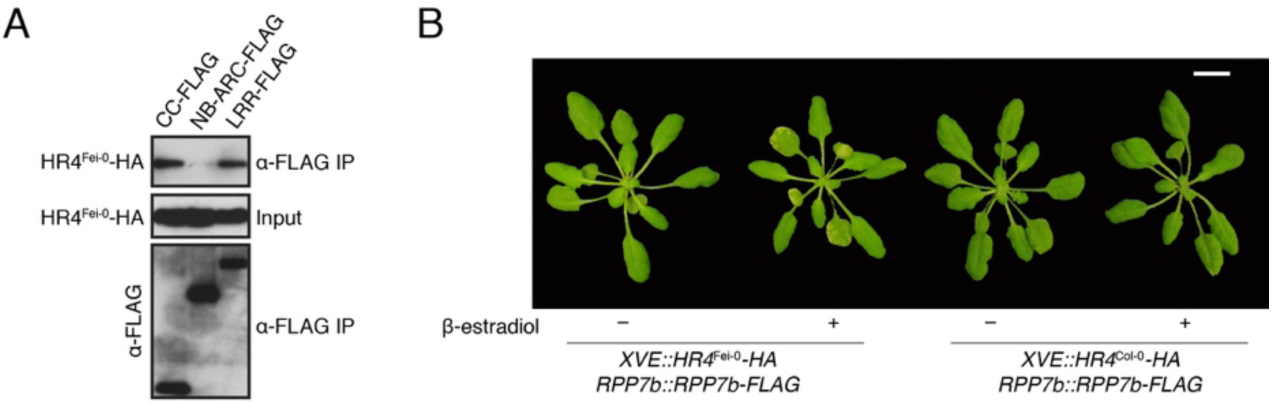
Association of HR4^Fei-0^ with RPP7b. (A) HR4^Fei-0^ interacts with CC and LRR domains of RPP7b in vivo, as shown by co-IP from *A. thaliana* protoplasts. (B) Transgenic *A. thaliana* plants used in Figure 2C, 2D and 2E, seven days after treatment with β-estradiol.

**Figure S3.**
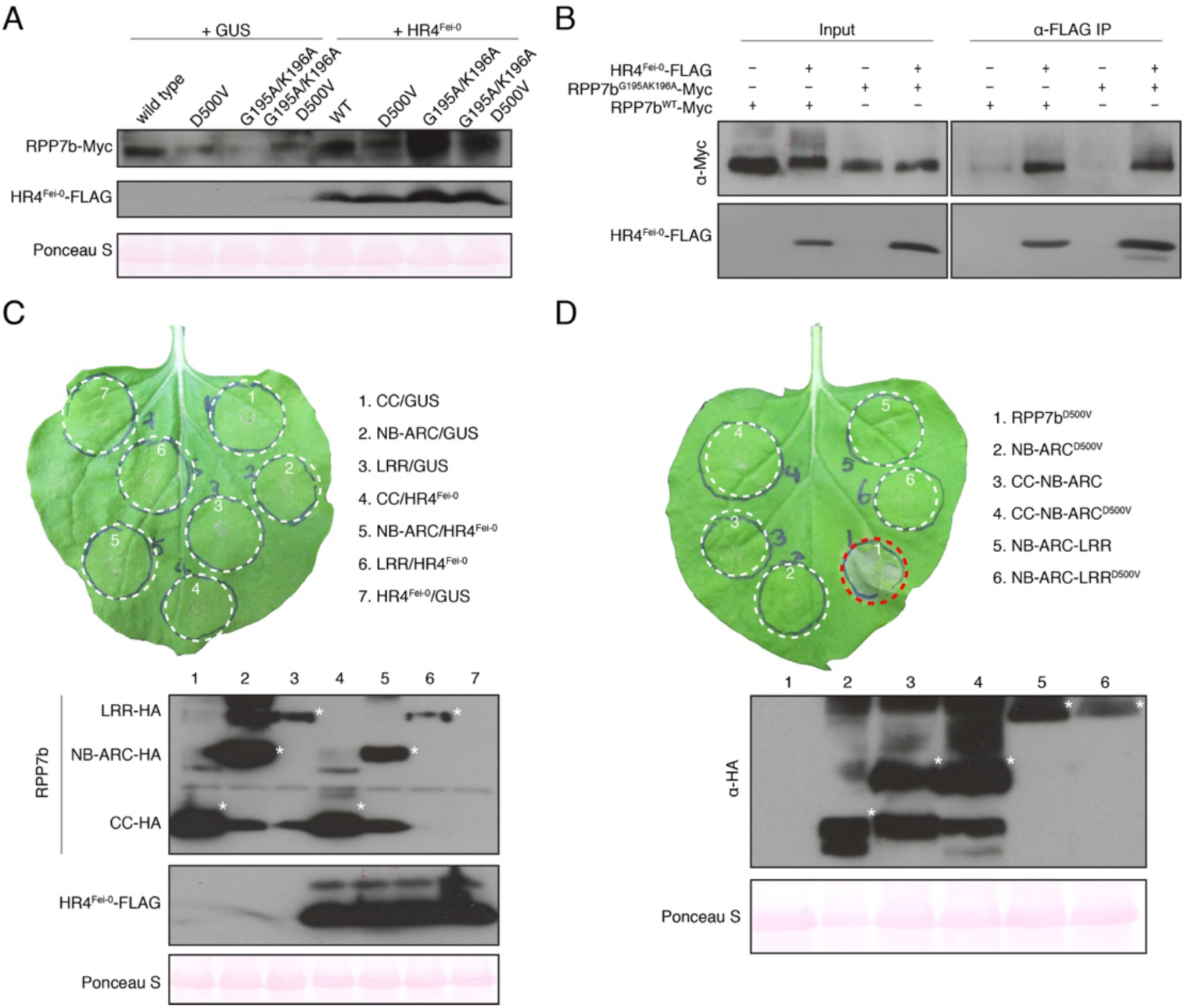
Protein accumulation of HR4^Fei-0^ and RPP7b variants. (A) Protein accumulation of P-loop and MHD motif mutants of RPP7b and of HR4^Fei-0^ in *N. benthamiana*, two days after *Agrobacterium* infiltration. Functional analyses shown in Figure 3C. (B) Interaction of HR4^Fei-0^ with P-loop mutant RPP7b variant, as shown by co-IP from *N. benthamiana*, two days after *Agrobacterium* infiltration. (C) Full-length RPP7b is required for activation of immune signaling. Top, lack of hypersensitive response induction by HA-tagged RPP7b domains in the presence or absence of HR4^Fei-0^-FLAG. Bottom, protein accumulation for experiments shown above. Asterisks indicate bands of the expected size. (D) MHD mutation in RPP7b NB-ARC domain is insufficient for hypersensitive response. Top, lack of hypersensitive response induction by RPP7b NB-ARC domain variants with MHD mutation D500V. RPP7b^D500V^ was used as a positive control. Bottom, protein accumulation for experiments shown above. Asterisks indicate the bands of the expected size. RPP7b^D500V^-HA, even though clearly phenotypically active, was undetectable, most likely because of rapid degradation after autoactivation.

**Figure S4.**
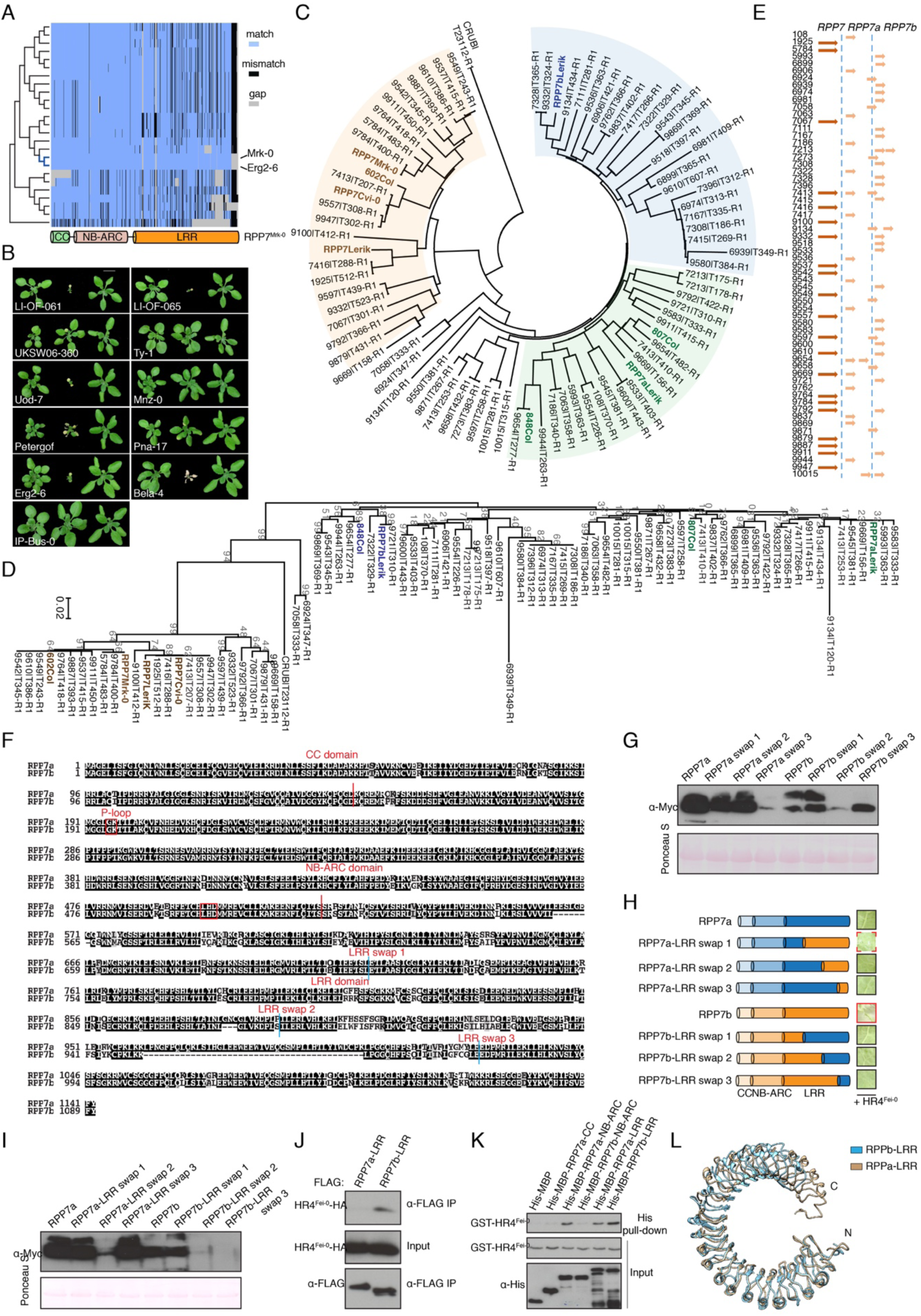
Requirement of RPP7 LRRs for HR4^Fei-0^-induced oligomerization. (A) Alignments of RPP7 related proteins from 64 accessions with RPP7^Mrk-0^ reveal sequence and structural polymorphisms, especially in the LRR domain. (B) Several accessions with *RPP7b*^Mrk-0^-like alleles produce hybrid necrosis when crossed to KZ10. F_1_ hybrids (middle) and different accessions (left) grown at 16°C. Some of the KZ10 control plants shown (always on the right) are identical, because several F_1_s were grown at the same time. Size bar corresponds to 1 cm. (C) Phylogeny of RPP7 full-length protein sequences from 64 accessions. Brown, RPP7^Col-0/Mrk-0^ clade; green, RPP7a^Lerik1-3^ clade; blue, RPP7b^Lerik1-3^ clade. (D) Phylogeny of RPP7 NB-ARC domain sequences from 64 accessions. Bootstrap support out of 100 given in grey. (E) RPP7 copy number variation in different accessions. RPP7a and RPP7b subclades are based on the phylogeny in (A). (F) Protein sequence alignment between RPP7a and RPP7b to indicate domain boundaries used for domain swaps. (G) Protein accumulation as determined by immunoblot for experiments shown in Figure 4A. (H) Hypersensitive response (red frames) induced by co-expression of HR4^Fei-0^ and domain swaps between RPP7a and RPP7b LRR domains in *N. benthamiana*, seven days after *Agrobacterium* infiltration. Experiments were performed at least twice. (I) Protein accumulation as determined by immunoblot for experiments in (H). (J) HR4^Fei-0^ interacts more effectively with the RPP7b^LRR^ than the RPP7a^LRR^ domain, as shown by co-IP from *A. thaliana* protoplasts. (K) HR4^Fei-0^ interacts more strongly with the RPP7b^LRR^ than the RPP7a^LRR^ domain in vitro, as shown by pull-down assays with proteins purified from *E. coli*. (L) Superposition of homology models of RPP7a and RPP7b LRR structures.

**Figure S5.**
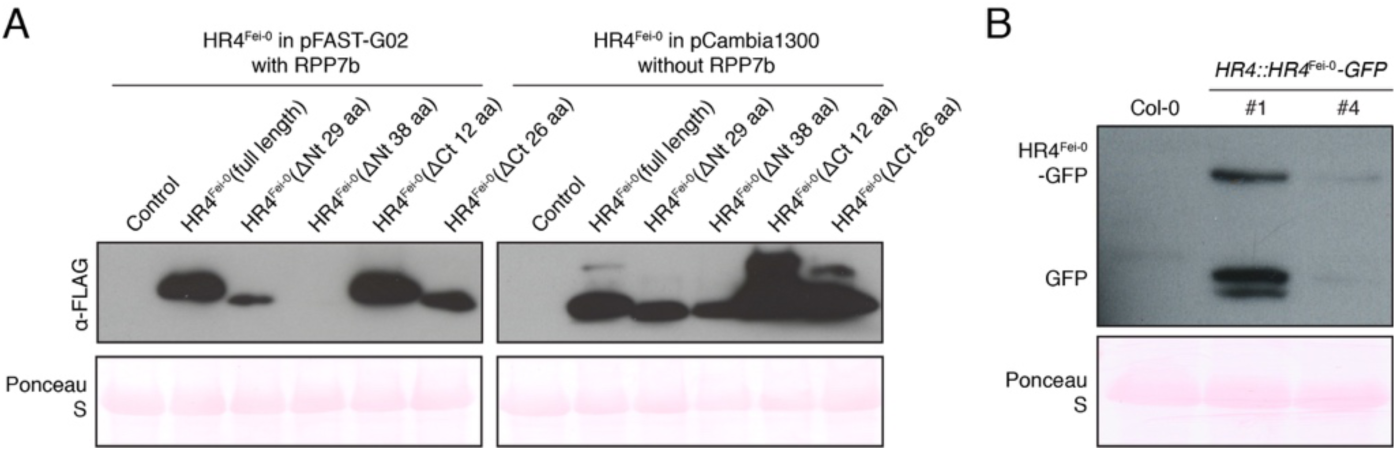
RPP7b-independent activity of HR4^Fei-0^. (A) Protein accumulation as determined by immunoblot for experiments in Figure 5A. (B) Protein accumulation as determined by immunoblot for transgenic *A. thaliana* lines in Figure 5B.

**Figure S6.**
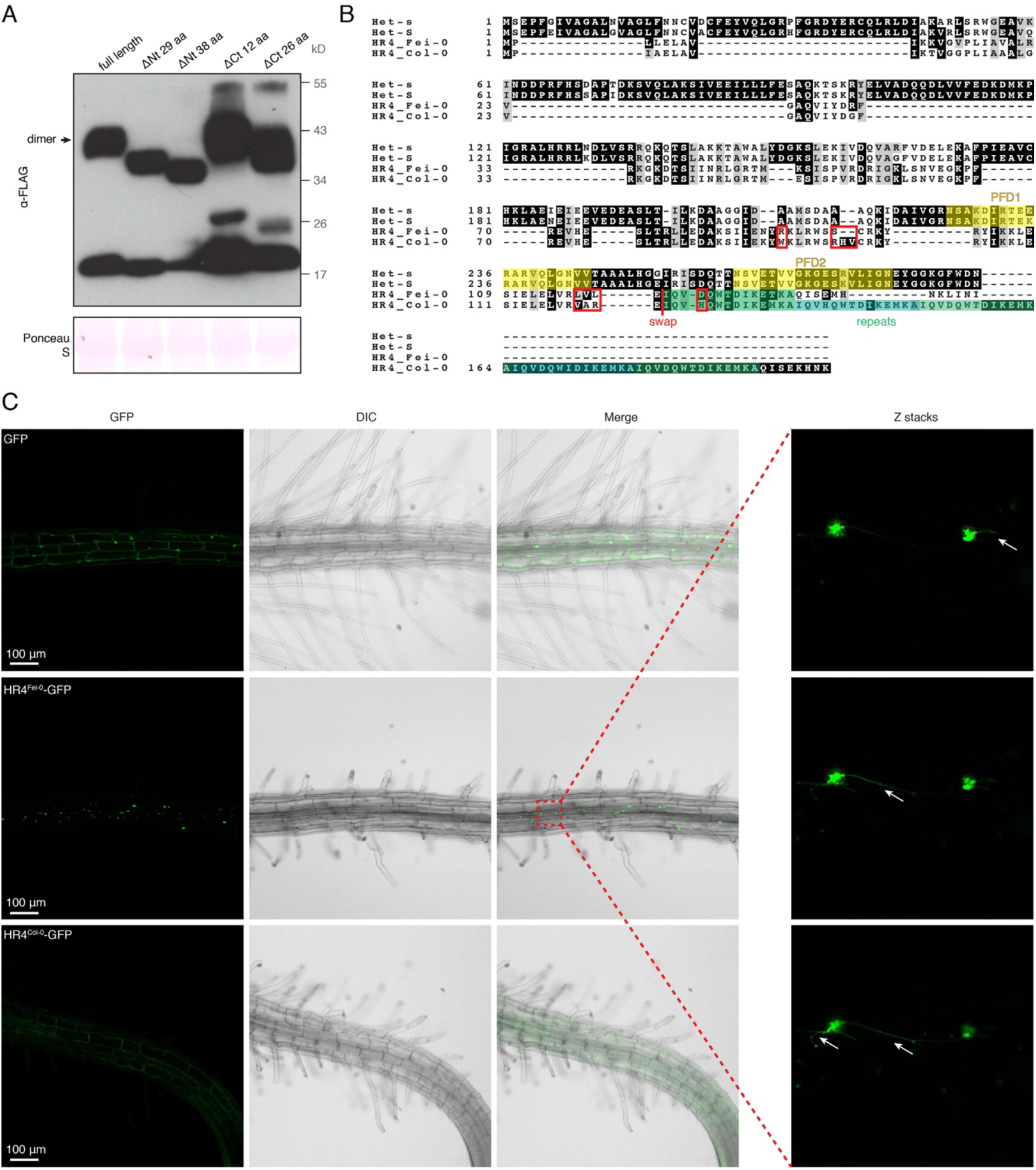
Aggregation of HR4^Fei-0^ contributes to its cytotoxic activity. (A) Protein accumulation for truncated HR4^Fei-0^ variants expressed in *N. benthamiana*; arrow indicates SDS-resistant dimers. (B) Protein sequence alignment between fungal Het-s, Het-S, HR4^Fei-0^ and HR4^Col-0^, indicating Het-s/Het-S prion-forming domain (PFD, yellow), HR4 repeats (green), and HR4 domain boundary used for HR4^Col-0^/HR4^Fei-0^ swaps. (C) Confocal images of transgenic lines expressing inducible GFP-only, HR4^Fei-0^-GFP, and HR4^Col-0^-GFP after 10 h treatment with β-estradiol. Arrows point to Z-membrane like structures. DIC, Differential Interference Contrast.

